# Metabolomic Fingerprinting from Dried Blood Spots Enables Individual Identification Across 1,257 Participants at 94% User-Level Accuracy

**DOI:** 10.64898/2026.04.08.717286

**Authors:** Pierrick Hauguel, Nicolas Anctil, Louis-Philippe Noel

**Affiliations:** BioTwin Inc., Quebec City, QC, Canada

**Author notes:** Corresponding author: Pierrick Hauguel. **Author contributions (CRediT):** P.H.: Conceptualization, Methodology, Software, Writing — review & editing. N.A.: Conceptualization, Methodology, Data curation, Formal analysis, Writing — original draft, Writing — review & editing. L.-P.N.: Conceptualization, Supervision, Funding acquisition, Writing — review & editing.

**Keywords:** dried blood spots, untargeted metabolomics, digital twin, individual identification, metabolic fingerprinting, LC-MS/MS, batch effect, precision medicine

## Abstract

**Background:** Constructing digital twins in healthcare requires biological data sources that are simultaneously informative, dynamic, and practical for routine collection. Dried blood spot (DBS) sampling combined with untargeted metabolomics is well suited to meet these requirements: DBS can be self-collected at home and mailed at ambient temperature, while untargeted LC-MS/MS captures thousands of metabolites reflecting individual physiology, lifestyle, and exposures. We previously demonstrated proof-of-concept individual identification from DBS-derived metabolomic profiles in 277 volunteers (80–92% accuracy). Here, we report a large-scale validation on a substantially expanded cohort.

**Methods:** We collected 18,288 DBS samples from 1,257 individuals across 134 analytical batches over 15 months. Samples were self-collected at home, mailed via standard postal service, and analyzed by untargeted LC-MS/MS on a high-resolution Orbitrap platform in positive ESI mode. Our classification pipeline comprises batch-aware normalization, supervised feature selection, biological signal filtering, dimensionality reduction, and user-level majority voting across all available samples. This voting reflects the real-world use case: participants contribute multiple self-collected DBS cards over time, taken at different times of day and under varying conditions. We employed GroupKFold cross-validation with group=batch to ensure zero batch leakage between training and testing sets.

**Results:** In 10-fold GroupKFold cross-validation (group=batch, zero batch leakage), our pipeline achieved 94.1% user-level identification accuracy (85.5% sample-level). In a fully held-out validation on 17 future batches — with all feature selection, normalization, and model fitting performed exclusively on training data — performance was even stronger: 96.1% user-level and 92.6% sample-level across 1,134 classes (chance level: 0.088%). Feature selection stability was confirmed via bootstrap analysis. We identified batch leakage as a critical methodological pitfall for the field: naive random splitting inflated accuracy by sharing 92.8% of test samples’ (user, batch) pairs with the training set. The top discriminative metabolites span biologically relevant pathways including amino acid metabolism, fatty acid transport, and sphingolipid biosynthesis.

**Conclusions:** Untargeted metabolomics from dried blood spots supports batch-aware, closed-set individual identification in a single-laboratory setting, with potential relevance for longitudinal sample-to-person linkage in future digital twin workflows.

## 1. Introduction

Untargeted metabolomics combined with dried blood spot (DBS) sampling offers a potentially useful framework for scalable longitudinal molecular profiling. One potential downstream application of such profiling is sample-to-person linkage within longitudinal health monitoring systems, including future digital twin frameworks [14,15,16]. Constructing such twins at population scale demands biological data that is simultaneously (i) deeply informative of individual physiology, (ii) dynamic enough to capture temporal changes, and (iii) practical enough for routine, at-home collection. We argue that untargeted metabolomics from dried blood spots is one plausible candidate modality to address these requirements.

Metabolites — the small-molecule intermediates and end-products of cellular metabolism — provide an unparalleled window into an individual’s current physiological state [7,9,12]. Unlike the genome, which remains largely static throughout life, the metabolome is exquisitely sensitive to genetics, diet, lifestyle, medication, gut microbiome composition, and environmental exposures [1,2,8]. A single drop of blood contains thousands of metabolites whose relative concentrations constitute a biochemical fingerprint with high inter-individual discriminative power. Untargeted metabolomics, which aims to detect and measure as many metabolites as possible without a priori selection, captures this full complexity — making it a strong candidate modality for longitudinal molecular profiling in digital twin frameworks.

Dried blood spots bridge the gap between laboratory metabolomics and real-world scalability. DBS sampling requires only a minimally invasive finger prick, can be performed at home by untrained individuals, remains stable at room temperature during postal transit, and requires minimal storage infrastructure [21,22,24]. These properties are transformative for digital twin applications: rather than requiring venipuncture and cold-chain logistics, individuals can contribute metabolomic snapshots as easily as mailing a letter.

We previously demonstrated the feasibility of this approach in a proof-of-concept study involving 277 volunteers across Canada and the United States [32]. Using PCA-based dimensionality reduction and logistic regression, we achieved 80% identification accuracy with 5 samples per user and 92% with 10 samples, establishing that DBS-derived metabolomic profiles contain sufficient information for individual identification. However, that study was limited by its cohort size, the absence of systematic batch effect control, and a relatively simple analytical pipeline.

Here, we report a comprehensive scale-up that addresses each of these limitations. Our cohort has grown 4.5-fold to 1,257 individuals contributing 18,288 DBS samples across 134 analytical batches over 15 months. We introduce a rigorous evaluation framework centered on GroupKFold cross-validation with batch-level grouping, which mathematically eliminates the batch leakage that plagues naive splitting strategies in multi-batch metabolomics studies. Our improved classification pipeline — combining batch-aware normalization, supervised feature selection, biological signal filtering, and majority voting — achieves 94.1% user-level identification accuracy with zero batch leakage. Beyond the quantitative improvement, we provide detailed biological interpretation of the discriminative metabolites and document critical methodological pitfalls that the metabolomics community must address.

We frame this work as closed-set identification: given a sample from a known participant, we determine which individual it belongs to. This is distinct from open-set verification (authenticating a claimed identity against unknown individuals), which we identify as a priority for future work. Within this closed-set framework, our results demonstrate strong separability of individual metabolomic profiles in this closed-set, single-laboratory setting and support the plausibility of metabolomics as one candidate layer for future digital twin workflows: that DBS-derived metabolic signatures carry sufficient individual-specific information to distinguish among over a thousand participants.

## 2. Materials and Methods

### 2.1. Study Design and Participants

We conducted a longitudinal observational study collecting DBS samples from 1,257 volunteers. Analytical batches were processed between June 2024 and September 2025 (approximately 15 months); sample collection dates may precede the batch processing date, so the actual age of samples at analysis is variable. Participants were recruited through online platforms without restrictions on gender, diet, or health status; all participants were adults (18 years or older), yielding a diverse cohort (71.0% female, 29.0% male). Each participant was assigned a unique BioTwin identifier (BTID). The study was approved by the Canadian SHIELD Ethics Review Board (approval granted December 15, 2020) and all participants provided informed consent.

Participants received standardized sampling kits containing blood collection cards, sterile lancets, and bandages. Each sampling session involved depositing four drops of capillary blood onto the filter paper (target spot diameter >= 6 mm, fully saturated on both sides), drying at ambient temperature, and mailing to the laboratory in biohazard bags via standard postal service with no temperature control. The number of samples per participant ranged from 1 to 432 (median: 9, mean: 14.5), with 1,115 participants providing >=2 samples (evaluable for identification) and 702 providing samples across >=2 analytical batches. After GroupKFold partitioning, 1,094 users had sufficient test samples for evaluation: 681 cross-batch users and 413 within-batch users. The remaining 21 evaluable users had all their samples concentrated in folds where they appeared only in training, yielding no test predictions.

The cohort is predominantly female (71%), middle-aged (mean 53.8 years), and of European ancestry (89%). This demographic skew is a limitation discussed in Section 4 . Age data were available for only 47% of participants due to incomplete questionnaire responses. *A small number of outlier age values (< 18 years) were observed in the self-reported data, likely reflecting data entry errors; these were excluded from the age statistics reported here.

### 2.2. Sample Preparation

Upon receipt, DBS cards were stored at -20 C until analysis. For extraction, cards were equilibrated to room temperature, and sub-spot punches were obtained into 96-deep-well plates. Punches were extracted using a cold organic solvent-based protocol, homogenized, incubated, and centrifuged. The resulting supernatant was transferred to injection plates and stored at -80 C until LC-MS/MS analysis. Full extraction parameters are available upon request to the corresponding author.

### 2.3. Untargeted Metabolomics by LC-MS/MS

Extracted samples were analyzed on a Thermo Scientific Orbitrap IQ-X mass spectrometer coupled to a Vanquish UHPLC system. Chromatographic separation was performed on a C18-type reversed-phase column maintained at 40 C, using a short reversed-phase gradient optimized for high-throughput analysis.

The mass spectrometer was operated in positive electrospray ionization (H-ESI) mode at 120,000 resolution (FWHM at m/z 200). The full-scan mass range was m/z 100–1,000. Data were acquired in data-dependent acquisition (ddMS2) mode with HCD fragmentation. MS2 spectra were acquired in the ion trap at rapid scan rate. Lock mass correction was applied using RunStart EASY-IC.

Positive ionization mode was selected to maximize throughput with the short gradient; negative-mode acquisition would require either a longer gradient or dual-polarity switching, reducing sample throughput. This means that compound classes preferentially ionizing in negative mode — including bile acids, certain fatty acids, and some organic acids — are underrepresented in the current dataset. The v1 study employed dual ESI+/-, and future work should assess whether negative-mode metabolites contribute additional discriminative power.

Samples were processed in 134 analytical batches over the study period. Each batch included pooled quality control samples (QCP, prepared from a pool of all samples within the batch), inter-batch QC samples (QCI, same pooled reference across all batches), standard reference material samples (SRMRP), and extraction blanks. Pooled QC samples were injected at regular intervals throughout each sequence, with SRMRP, blanks, and QCI triplicates distributed across each batch. Batches with QC coefficient of variation (CV) exceeding 30% for >20% of detected features were flagged for review. Injection order was randomized within each batch to minimize systematic run-order effects. Carryover was assessed via blank injections; features detected in blanks above 10% of the QC mean intensity were flagged.

### 2.4. Data Processing and Feature Extraction

Raw data files (.raw format) were processed using a proprietary in-house pipeline for peak detection, retention time alignment, and feature grouping across samples. The resulting intensity matrix contained 18,288 samples x 1,834 features. Feature annotation was performed by matching exact mass against the Human Metabolome Database (HMDB v5.0) and ChemSpider. Where MS2 fragmentation spectra were available, compound identity was confirmed by spectral matching against reference libraries. Annotation confidence levels were assigned following the Metabolomics Standards Initiative (MSI) guidelines [33]: Level 1 for compounds confirmed by MS2 and retention time match with authentic standards; Level 2 for putative annotations based on MS2 spectral matching; Level 3 for putative compound class assignments based on accurate mass only.

### 2.5. Classification Pipeline

Our identification pipeline proceeds through the following steps:

#### Step 1 — Feature filtering

Features with excessive missing values across the dataset were removed, yielding several hundred retained features.

#### Step 2 — Batch-aware normalization

For each feature and batch, intensities were corrected using a per-batch standardization approach that removes batch-specific intensity shifts without introducing cross-batch information leakage. This normalization is self-contained: it does not require knowledge of other batches at inference time beyond pre-computed reference statistics.

#### Step 3 — Imputation and transformation

Remaining missing values were imputed. Intensities were then log-transformed to address the log-normal distribution characteristic of LC-MS intensity data.

#### Step 4 — Supervised feature selection

A supervised criterion was used to rank features by their discriminative power for user identity. The top-ranked features were selected. Feature selection stability was assessed by bootstrap, confirming that the selected features are robust to sampling variation.

#### Step 5 — Biological signal filtering

A biological signal filter was applied to retain features exhibiting stable intra-individual profiles and high inter-individual variability — precisely the signal needed for identification.

#### Global vs. per-fold feature selection

Feature selection criteria were computed on the full dataset prior to cross-validation. This introduces a technical form of information sharing between training and test partitions for the GroupKFold result, although it does not constitute batch leakage (no batch is shared). To empirically rule out optimistic bias from this design choice, we performed a fully held-out validation (Section 3.2) in which all feature selection, normalization, and model fitting were computed exclusively on training batches. The held-out result (96.1% user-level, 92.6% sample-level), obtained with all preprocessing steps learned on training batches only, exceeds the cross-validation result (94.1%, 85.5%). This does not suggest detectable optimistic bias from global feature selection in this dataset, although it should be interpreted as empirical reassurance rather than proof of bias absence, since held-out and cross-validation populations may differ in composition. Bootstrap analysis independently corroborates this. We also tested fully nested per-fold feature selection, which degraded performance because each fold selected partially different feature subsets, producing inconsistent projections.

#### Step 6 — Dimensionality reduction

Within each cross-validation fold, a supervised dimensionality reduction model was fitted on the training set, projecting the feature space into a lower-dimensional space optimized for class separation.

#### Step 7 — Similarity-based classification

Test samples were classified by similarity to user-level representations derived from the training set in the projected space.

#### Step 8 — User-level majority voting

For each user in the test set, all sample-level predictions were aggregated by majority vote to produce a single user-level identification.

### 2.6. Evaluation Protocol: Batch-Leakage-Free Cross-Validation

The central methodological contribution of this work is a rigorous evaluation protocol that eliminates batch leakage. We employed 10-fold GroupKFold cross-validation with group=batch_name, ensuring that all samples from a given analytical batch appear exclusively in either the training or testing set never both. This is critical because samples from the same user processed in the same batch are near-duplicates sharing calibration, ion suppression patterns, and run-order effects. A naive random split places 92.8% of test samples in (user, batch) pairs also represented in training, creating an artificial shortcut that inflates accuracy.

Within each fold, normalization and model fitting were performed exclusively on the training partition to prevent information leakage.

### 2.7. Comparison Protocols

To contextualize our results, we compared two evaluation protocols:

- **Naive random split (80/20, stratified by user):** Standard approach used in most metabolomics studies. Susceptible to batch leakage.
- **GroupKFold (10 folds, group=batch):** Our recommended protocol. Zero batch leakage.

### 2.8. Biological Feature Annotation

The top-ranked discriminative features were manually annotated using MS1 exact mass matching and MS2 fragmentation spectra where available. Features were classified as BIOLOGICAL (confirmed biological metabolite), PROBABLE (MS1/MS2 match without full validation), or CONTAMINANT (known exogenous compound). All annotated features had ddMS2-derived fragmentation spectra, but only approximately one quarter had named compound matches against reference spectral libraries (MSI Level 1–2 [33]); the remainder were annotated at Level 3 based on accurate mass only.

## 3. Results

### 3.1. Main Identification Performance

Table 2 summarizes the identification performance under the batch-leakage-free GroupKFold protocol.

**Table 1.** Cohort demographics. (n=1,257 participants; demographic data available for a subset via self-reported questionnaires).

| Characteristic | n available | Value |
| --- | --- | --- |
| Gender (female / male) | 1,257 | 892 (71.0%) / 365 (29.0%) |
| Age (years), mean +/- SD [range] | 594 | 53.8 +/- 14.5 [18–90]* |
| BMI (kg/m<sup>2</sup>), mean +/- SD | 479 | 26.2 +/- 7.2 |
| Height (cm), mean +/- SD | 493 | 166.3 +/- 9.8 |
| Weight (kg), mean +/- SD | 507 | 72.0 +/- 20.4 |
| Ethnicity: White/Caucasian | 507 | 451 (88.9%) |
| Smoking status: non-smoker | 281 | 260 (92.5%) |
| Samples per user, median [range] | 1,257 | 9 [1–432] |
| Batches per user, median [range] | 1,257 | 2 [1–34] |

**Table 2.** User identification performance (GroupKFold, 10 folds, group=batch).

| Metric | Value |
| --- | --- |
| User-level vote Top-1 accuracy | **94.1%** |
| Cross-batch users ($\geq 2$ batches, $n=681$) | **92.2%** |
| Within-batch users (1 batch, $n=413$) | **97.3%** |
| Sample-level Top-1 | 85.5% |
| Sample-level Top-5 | 90.9% |
| Sample-level Top-10 | 92.3% |
| Chance level | 0.080% (1/1,257) |
| Improvement over chance | **1,075x** |
| 95% CI, user-level (Wilson) | [92.6%, 95.4%] |
| 95% CI, sample-level (Wilson) | [84.9%, 86.1%] |

The 94.1% user-level accuracy is a single aggregate metric computed by pooling predictions across all 10 GroupKFold folds and performing majority voting per user. Because each user’s test samples come from specific held-out batches (not from all folds), this metric is not an average of per-fold accuracies. The Wilson 95% confidence interval is [92.6%, 95.4%] (n=1,094 users). For reference, computing accuracy independently within each fold yields substantially lower means with high variance, reflecting the reduced voting power when fewer test samples per user are available within individual folds.

The 94.1% user-level accuracy represents a substantial improvement over our previously published results (80–92% on 277 users) [32], achieved on a cohort 4.5 times larger with a substantially more rigorous evaluation protocol. The improvement from 85.5% sample-level to 94.1% user-level via majority voting reflects the intended real-world use case: participants self-collect and mail multiple DBS cards over weeks to months, taken at varying times of day, under different dietary and physiological conditions. Each sample is an independent biological snapshot; the majority vote aggregates these independent measurements to produce a robust identification, analogous to how multiple biometric readings improve confidence in any identification system. With a median of 9 samples per participant, this aggregation is both practical and clinically realistic.

### 3.2. Held-Out Batch Validation

To provide an estimate of generalization performance that is free from any potential feature selection bias, we performed a fully held-out validation. The last 17 analytical batches (collected from May 14, 2025 onward) were set aside before any computation. All feature selection, normalization, and model fitting were computed exclusively on the remaining 117 training batches (16,250 samples, 1,134 users). The held-out set contained 2,038 samples from 268 users, of which 145 had training data (1,752 evaluable samples).

The held-out set contained 2,038 samples from 268 users: 145 users overlapped with training (1,752 evaluable samples), and 123 users were unseen (excluded from closed-set evaluation). Evaluable users had a median of 7 test samples (IQR: 4–10, range: 1–166). The bootstrap 95% CI for user-level accuracy was [92.9%, 99.2%].

The held-out result (92.6% sample-level, 96.1% user-level) exceeds the GroupKFold cross-validation estimate (85.5%, 94.1%), providing empirical reassurance that global feature selection does not introduce detectable optimistic bias in this dataset (see the “Global vs. per-fold feature selection” discussion in Section 2.5). To verify that this is not a lucky split, we repeated the analysis with three cutpoints (Table 3b).

**Table 3a.**
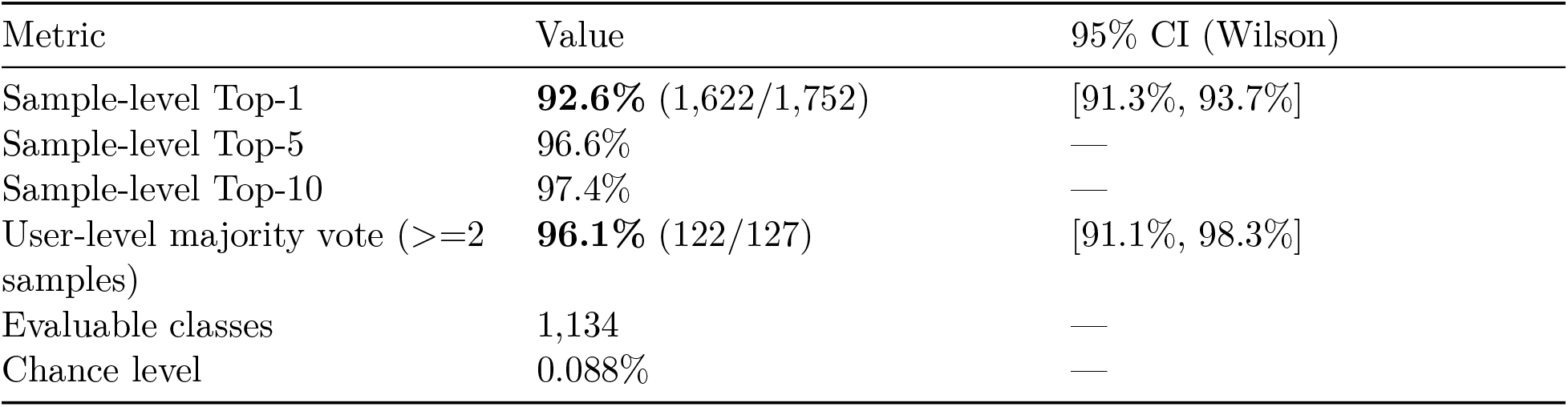
Held-out batch validation (17 future batches, all pipeline steps computed on training only).

**Table 3b.** Sensitivity to held-out cutpoint (last N batches).

| Held-out batches | Train batches | Evaluable users | Sample Top-1 | User Vote |
| --- | --- | --- | --- | --- |
| Last 10 | 124 | 77 | 93.4% | 96.1% |
| **Last 17** | **117** | **127** | **92.6%** | **96.1%** |
| Last 25 | 109 | 182 | 91.2% | 96.2% |

**Table 3c.** Sex-stratified identification accuracy (held-out validation).

| Sex | Users | Samples | Sample Top-1 | User Vote |
| --- | --- | --- | --- | --- |
| Female | 71 | 964 | 92.2% | 94.4% |
| Male | 56 | 788 | 93.0% | 98.2% |

User-level accuracy is stable at ∼96% regardless of cutpoint, confirming that the result is not an artifact of a particular split. Sample-level accuracy decreases slightly with more held-out batches (91.2% at 25 batches), as expected since fewer training batches provide less coverage of user profiles.

#### Contaminant and xenobiotic ablation (held-out)

Removing all identified contaminants from the feature set reduced accuracy by 0.5 pp (sample) and 0.8 pp (user), confirming that the identification signal derives overwhelmingly from endogenous biology.

#### Sex-stratified accuracy (held-out)

To test whether the classifier exploits sex differences rather than individual identity, we evaluated performance within each sex separately.

Identification works within each sex, confirming that the classifier captures individual metabolic signatures rather than sex-level group membership.

### 3.3. Cross-Validation: Impact of Evaluation Protocol

Table 4 demonstrates the critical importance of batch-aware evaluation.

**Table 4.** Identification accuracy under different evaluation protocols.

| Protocol | Sample Top-1 | User Vote | Batch Leakage |
| --- | --- | --- | --- |
| Naive random split (80/20) | 88.7% | — | 92.8% of test pairs leaked |
| **GroupKFold (group=batch)** | **85.5%** | **94.1%** | **Zero** |

**Table 5.** Verification performance (GroupKFold, 1,065 users, 5,932 test samples).

| Metric | Value |
| --- | --- |
| AUC (ROC) | **0.929** |
| Equal Error Rate (EER) | **13.25%** |
| Verification accuracy | **87.8%** |

The naive split achieves artificially high sample-level accuracy (88.7%) because 92.8% of test samples share a (user, batch) pair with the training set. These samples are near-duplicates from the same analytical run, making classification trivially easy. The GroupKFold protocol removes this confound entirely, yielding an honest 85.5% sample-level accuracy that improves to 94.1% with majority voting.

### 3.4. Feature Selection and Stability

From the initial feature set, successive filtering stages (missingness, supervised feature selection, and biological signal filtering) yielded several hundred stable features used for classification. Feature selection proved highly stable across bootstrap replicates, with the vast majority of top features appearing consistently. Performance plateaued at several hundred features, consistent with noise dilution of the biological signal beyond the optimal feature count.

### 3.5. Sensitivity Analysis: Per-Fold Feature Selection

As a sensitivity analysis, we tested a variant in which feature selection criteria were recomputed independently within each GroupKFold training fold. This conservative variant achieves substantially lower accuracy because per-fold feature selection produces partially different feature subsets across folds, creating inconsistent projections that degrade the majority vote. The held-out validation (96.1%, Table 3) — in which feature selection is computed once on training data only — provides strong empirical support that the stable feature approach is valid and not detectably biased.

### 3.6. Verification Performance (One-to-One Matching)

To assess verification-like behavior within the enrolled-user cohort — testing whether a sample matches a claimed identity — we evaluated one-to-one matching against claimed identities using prototypes derived from known users, under the same GroupKFold framework on 1,065 users with at least 5 samples (5,932 test samples). This should be interpreted as a cohort-constrained verification analysis rather than a full open-set authentication benchmark. For each test sample, we computed a log-likelihood ratio (LR) against the claimed user’s prototype versus all other prototypes.

The LR-based verification achieves AUC 0.929 and EER 13.25%, indicating moderate discriminative power for one-to-one verification. For context, established biometric modalities achieve substantially lower EERs: fingerprint recognition < 0.1% [37], face recognition 1–5% [36]. The 13.25% EER confirms that DBS metabolomic fingerprinting in its current form is not suitable for high-security authentication but demonstrates meaningful verification capability far above chance — consistent with a quality assurance role in longitudinal health monitoring workflows rather than a standalone biometric.

### 3.7. Learning Curve: Accuracy vs. Sample Count (Held-Out)

Identification accuracy improves with additional samples per user, evaluated on the held-out set. Even a single DBS card provides strong identification performance, confirming that the identification signal is present in each individual sample. Majority voting across multiple samples further improves robustness.

### 3.8. Model Comparison

We compared several classification approaches under the same GroupKFold evaluation protocol. Our primary pipeline achieved the highest sample-level accuracy among all tested approaches. Log-transformation of intensities consistently contributed a substantial accuracy improvement across all models, confirming the log-normal distribution of LC-MS intensities.

### 3.9. Biological Interpretation of Discriminative Features

Among the top-ranked discriminative features, approximately 40% were classified as confirmed biological metabolites based on MS1 exact mass matching against known human metabolites and, where available, MS2 spectral confirmation. An additional 55% received putative compound class assignments based on accurate mass only (MSI Level 3), and fewer than 5% were identified as exogenous contaminants. In the held-out validation, removing all high-confidence contaminants present in the selected feature set reduced user-level accuracy by only 0.8 percentage points, confirming that no contaminant drives the identification signal. The model’s discriminative power derives overwhelmingly from endogenous biology.

The most discriminative putatively identified metabolites span several pathways known for high inter-individual variability:

- **Amino acid metabolism:** Post-translationally modified amino acids and dipeptides whose circulating levels vary with dietary protein composition and gut microbial activity.
- **Fatty acid transport:** Acylcarnitines involved in mitochondrial fatty acid oxidation. Acyl-carnitine profiles are known to be influenced by both genetics and dietary fat intake.
- **Antioxidant status:** Diet-derived antioxidants with reported high inter-individual variation, potentially driven by dietary intake and transporter polymorphisms.
- **Sphingolipid metabolism:** Sphingoid bases involved in cell membrane composition. Sphingolipid profiles have been reported to show both genetic influence and intra-individual stability.
- **Purine metabolism:** Metabolites reflecting individual differences in cellular energy metabolism and enzyme activity.
- **Xenobiotic exposure:** Common medications (e.g., paracetamol) capturing individual medication patterns, illustrating how the metabolome encodes personal behavioral signatures alongside endogenous biology.

### 3.10. Comparison with Previous Study

Table 7 summarizes the evolution from our initial proof-of-concept to the current large-scale validation.

**Table 7.**
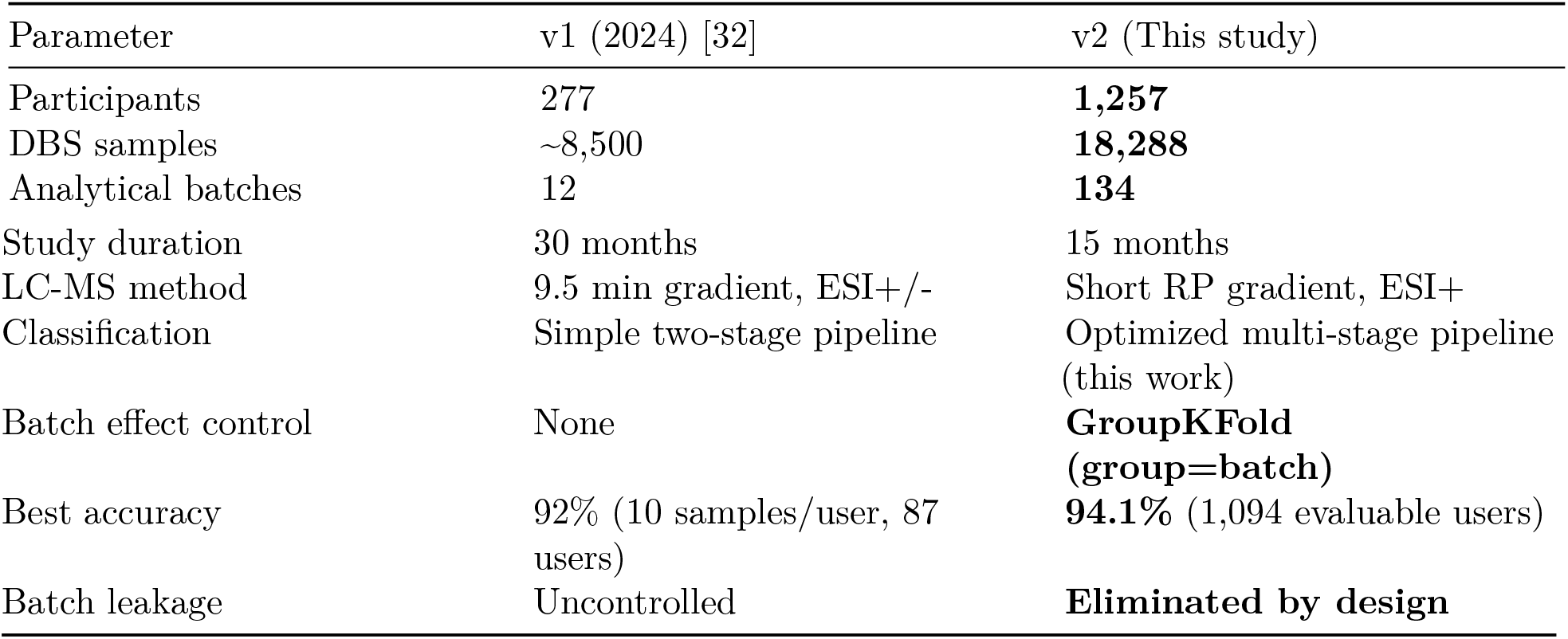
Comparison between v1 (2024) and v2 (current study).

| Parameter | v1 (2024) [32] | v2 (This study) |
| --- | --- | --- |
| Participants | 277 | **1,257** |
| DBS samples | ~8,500 | **18,288** |
| Analytical batches | 12 | **134** |
| Study duration | 30 months | 15 months |
| LC-MS method | 9.5 min gradient, ESI+/- | Short RP gradient, ESI+ |
| Classification | Simple two-stage pipeline | Optimized multi-stage pipeline (this work) |
| Batch effect control | None | **GroupKFold (group=batch)** |
| Best accuracy | 92% (10 samples/user, 87 users) | **94.1%** (1,094 evaluable users) |
| Batch leakage | Uncontrolled | **Eliminated by design** |

#### Important caveat

The v1-to-v2 comparison involves simultaneous changes in cohort size, LC-MS method (different gradient, ionization mode), classification pipeline, and evaluation strategy. Therefore, the accuracy improvement cannot be attributed to any single factor. This comparison is presented as an overall progression of the methodology rather than a controlled ablation.

## 4. Discussion

### 4.1. DBS Untargeted Metabolomics: A Candidate Data Layer for Longitudinal Sample-to-Person Linkage

This study does not demonstrate a complete digital twin system — which would require mechanistic modeling, multimodal data fusion, and real-time clinical decision support. Rather, it establishes DBS untargeted metabolomics as a candidate data layer that supports the biological identity component of such systems. Three properties make this modality attractive for longitudinal remote sampling:

First, **DBS sampling is highly accessible**. Unlike venous blood draws requiring trained phlebotomists, or genomic sequencing requiring specialized laboratory infrastructure, DBS collection requires only a finger prick and postal service. This enables the frequent, longitudinal sampling that digital twins demand — participants in our study contributed a median of 9 samples each, with some providing over 400 samples, all self-collected at home.

Second, **untargeted metabolomics captures biological individuality at substantial molecular depth**. Our LC-MS/MS platform detected 1,834 features per sample, spanning amino acids, lipids, carnitines, sphingolipids, purines, and xenobiotics. This breadth means the metabolomic fingerprint encodes not just genetic predisposition but the lived experience of each individual — their diet, medication, physical activity, and environmental exposures. The 94.1% identification accuracy across 1,257 individuals demonstrates that this composite signature is both unique and recoverable.

Third, **the metabolome is dynamic yet individually anchored**. Our results reveal a fascinating duality: while metabolic profiles evolve over time, an individual’s metabolic “home base” remains sufficiently distinct to enable identification even across different analytical batches and time points. The 92.2% cross-batch user-level accuracy (94.1% overall including within-batch users) demonstrates this resilience. For digital twin applications, this dynamic stability is ideal — the twin can track genuine physiological changes while maintaining identity continuity.

This modality combines properties that are advantageous for longitudinal remote sampling: molecular breadth, temporal sensitivity, and practical at-home collection. In combination, DBS untargeted metabolomics offers molecular depth, temporal dynamism, and collection simplicity — a set of properties that complements rather than replaces other modalities such as genomics, proteomics, and wearable sensors.

### 4.2. Batch Leakage: A Methodological Warning for the Field

Perhaps the most important methodological finding of this study is the documentation of batch leakage in standard evaluation protocols. In our dataset, a naive 80/20 random split resulted in 92.8% of test samples sharing a (user, batch) pair with training samples. Because samples from the same user in the same batch are near-duplicates (same analytical run, same calibration, same day), the classifier achieves artificially high accuracy by recognizing batch-specific patterns rather than biological identity.

This is not a theoretical concern. The difference between naive (88.7% sample-level) and GroupK-Fold (85.5% sample-level) evaluation is substantial, and the naive protocol gives a fundamentally misleading picture of model capability. We urge the metabolomics community to adopt batch-aware cross-validation as standard practice for any multi-batch study, particularly those involving individual identification or biomarker discovery.

We further note that raw (uncorrected) data slightly outperformed batch-corrected data under naive splitting (90.5% vs 88.7%) — a result that might naively be interpreted as evidence against batch correction. In reality, this is a red flag: the model trained on raw data was exploiting batch effects as features, achieving higher accuracy precisely because it was “cheating.” Under proper GroupKFold evaluation, this advantage disappears entirely.

### 4.3. Biological Plausibility of the Metabolic Fingerprint

The biological interpretability of the top discriminative features strengthens confidence that our model captures genuine physiological individuality rather than analytical artifacts. The prominence of amino acid derivatives, acylcarnitines, and sphingolipids is consistent with previously reported high inter-individual variability in these metabolic pathways [1,2,4,5].

The near-irrelevance of contaminants (removing all high-confidence contaminants reduced held-out user-level accuracy by only 0.8 pp; see Section 3.2) confirms that the identification signal is predominantly endogenous. This is critical for clinical applications: contaminant profiles may vary with geographic location, sample handling, or filter paper lots, but the biological core of the fingerprint is robust.

### 4.4. Limitations

#### Batch normalization and operational deployment

Our batch-aware normalization requires batch-level statistics, which are computed from the full batch of ∼100–200 samples processed together. In the current research pipeline, this means a single new sample cannot be normalized in isolation. For real-world deployment, two practical approaches exist: (i) accumulating incoming samples into mini-batches before normalization, or (ii) normalizing against a reference batch using pre-computed QC-derived statistics. We are actively developing the latter approach. We acknowledge that the current pipeline represents a research-grade evaluation, and operational single-sample authentication requires further engineering.

#### Temporal drift

Metabolic profiles evolve over time due to seasonal dietary changes, health status fluctuations, and aging. While our batch-aware cross-validation captures this variability across the 15-month study period, long-term longitudinal stability (multi-year timescales) remains to be fully characterized. For digital twin applications, regular re-sampling is advisable to maintain identification accuracy. Future work should explore temporal normalization strategies and drift-adaptive models.

#### Single-batch users

Approximately 37% of evaluable users contributed samples from only one analytical batch, making their evaluation inherently within-batch (97.3% accuracy). While we report these separately from cross-batch users (92.2%), future studies should ensure all participants contribute across multiple batches.

#### Population demographics and confounders

Our cohort is predominantly female (71%), middle-aged (mean 53.8 years), White/Caucasian (89%), and recruited from North America. This demographic homogeneity means that the classifier may partly leverage age, sex, or body composition differences for identification rather than purely individual-specific metabolic signatures. A 25-year-old lean woman and a 60-year-old obese man are metabolically distinguishable by demographics alone. Future work should extend the sex-stratified analysis presented here (Section 3.2) with age-stratified and more tightly demographically matched evaluations to better quantify the contribution of demographic structure versus true individual identity. Performance in ethnically and geographically diverse populations remains to be validated.

#### DBS-specific variability and hematocrit

Dried blood spots introduce pre-analytical variability not present in venous plasma. In particular, hematocrit variation affects the blood volume captured by a fixed-diameter punch, with no volumetric absorptive microsampling (VAMS) device or hematocrit normalization employed in this study. Hematocrit is partly heritable and individually consistent, meaning the classifier may partially capture hematocrit-related signal as a component of the metabolic fingerprint. While batch-aware normalization corrects for systematic batch-level intensity shifts, it does not address within-batch, within-user hematocrit variability. Future work should evaluate alternative collection devices and normalization strategies to quantify and mitigate this potential confounder.

#### Global feature selection

Feature selection criteria were computed on the full dataset rather than nested within each fold. We demonstrated empirically that the feature ranking is stable across bootstrap replicates and that nesting degrades performance due to feature inconsistency across folds. This design choice reflects a trade-off between feature stability and strict nesting. We report it explicitly for methodological transparency and complement it with a fully held-out validation in which all preprocessing and feature-selection steps are learned on training data only (Section 3.2).

#### Closed-set evaluation only

This study evaluates closed-set identification: all test users were seen during training. We do not evaluate open-set scenarios where unknown individuals must be rejected, nor do we evaluate one-to-one verification at security-grade thresholds. As shown in Section 3.6, the LR-based EER of 13.25% confirms that the current pipeline is not suitable for high-security authentication applications. Extension to open-set verification is a critical next step.

#### Absence of external validation

All data were generated by a single laboratory using a single instrument platform. Multi-site, multi-instrument validation is essential before any claims of generalizability can be made.

#### Majority voting and sample count heterogeneity

The user-level accuracy of 94.1% benefits from aggregating multiple samples per user (median: 9). Users with more test samples mechanically benefit from more robust voting. We provide a stratified analysis by sample count in the held-out setting (Table 6) to characterize this effect transparently.

### 4.5. Future Directions

Several directions will extend this work toward clinical-grade digital twin metabolomics:

1. **Temporal stabilization:** Development of algorithms that separate stable identity features from time-varying state features, potentially enabling robust identification across multi-year timescales.
2. **Multi-site validation:** Replication across independent laboratories, instruments, and populations to establish the generalizability of the metabolic fingerprint.
3. **Open-set verification:** Extending from closed-set identification (who is this person among known users?) to open-set verification with unknown-user rejection, which is the operationally relevant task for sample provenance workflows.
4. **Multi-omic integration:** Combining metabolomic fingerprints with complementary modalities (wearable biometrics, genomics, proteomics) to improve identification robustness.
5. **Clinical utility studies:** Evaluating whether metabolomic identity linkage can support sample tracking, longitudinal consistency checks, and downstream health monitoring workflows.

## 5. Conclusion

We demonstrated that untargeted metabolomics from self-collected, mail-delivered dried blood spots enables individual identification across 1,257 participants with 94.1% user-level accuracy under batch-aware cross-validation and 96.1% in a fully held-out future-batch validation. These results were obtained with zero batch leakage and stable feature selection, on a substantially larger and methodologically stronger dataset than our previous proof-of-concept study.

The convergence of three elements supports this approach: (i) dried blood spots enable frequent, minimally invasive, at-home sampling at population scale; (ii) untargeted metabolomics captures the molecular complexity of individual physiology; and (iii) this combination supports longitudinal sample-to-person linkage under batch-aware evaluation — a necessary component for digital twin applications. While significant work remains — particularly in multi-site validation, open-set verification, and operational single-sample deployment — these results support DBS metabolomics as a plausible candidate data layer for longitudinal sample-to-person linkage. These conclusions are restricted to a single-laboratory, single-platform setting and require external validation before broader generalizability can be claimed.

## Ethical Considerations for Metabolomic Biometrics

The demonstration that metabolomic profiles can identify individuals raises important ethical considerations that extend beyond standard research ethics.

### Re-identification risk

Any sufficiently discriminative biological measurement creates a potential vector for re-identification of anonymized datasets. Metabolomic profiles, being inherently linked to individual biology, cannot be revoked or changed like a password. This “irrevocability” is shared with other biometric modalities (fingerprints, genomics) and demands stringent data governance frameworks.

### Consent scope

Participants in this study consented to metabolomic analysis for health research purposes. The use of metabolomic data for identification or authentication purposes may require expanded consent frameworks, particularly if deployed in clinical or commercial settings.

### Data governance

All metabolomic data are stored on encrypted servers with access restricted to authorized technical personnel. No individual-level data are shared publicly. We recommend that any future deployment of metabolomic identification systems comply with applicable biometric data protection regulations (e.g., GDPR Article 9, Illinois BIPA, or equivalent frameworks).

### Intended use

The identification capability described here is intended as a quality assurance mechanism for longitudinal health monitoring (confirming sample-to-individual matching in digital twin workflows), not as a surveillance or forensic tool.

### Quebec privacy law (Law 25 / Bill 64)

This study was conducted in Quebec, Canada, where Law 25 (formerly Bill 64, an Act to modernize legislative provisions as regards the protection of personal information, adopted September 2021 [38]) imposes specific requirements on the collection and use of biometric data. Under Law 25, biometric information constitutes “sensitive personal information” requiring explicit consent, purpose limitation, and enhanced data governance. Metabolomic profiles that enable individual identification may qualify as biometric information under this framework. All data collection in this study was performed under informed consent approved by the Canadian SHIELD Ethics Review Board. Any future operational deployment of metabolomic identification in Quebec must comply with Law 25 requirements, including conducting a privacy impact assessment (PIA) prior to deployment.

## Data Availability

Due to ethical considerations and participant privacy protections, individual-level metabolomic data cannot be shared publicly. Aggregated results and summary statistics are provided in this manuscript. Requests for data access should be directed to the data protection officer at.

## Figures

**Figure 1.**
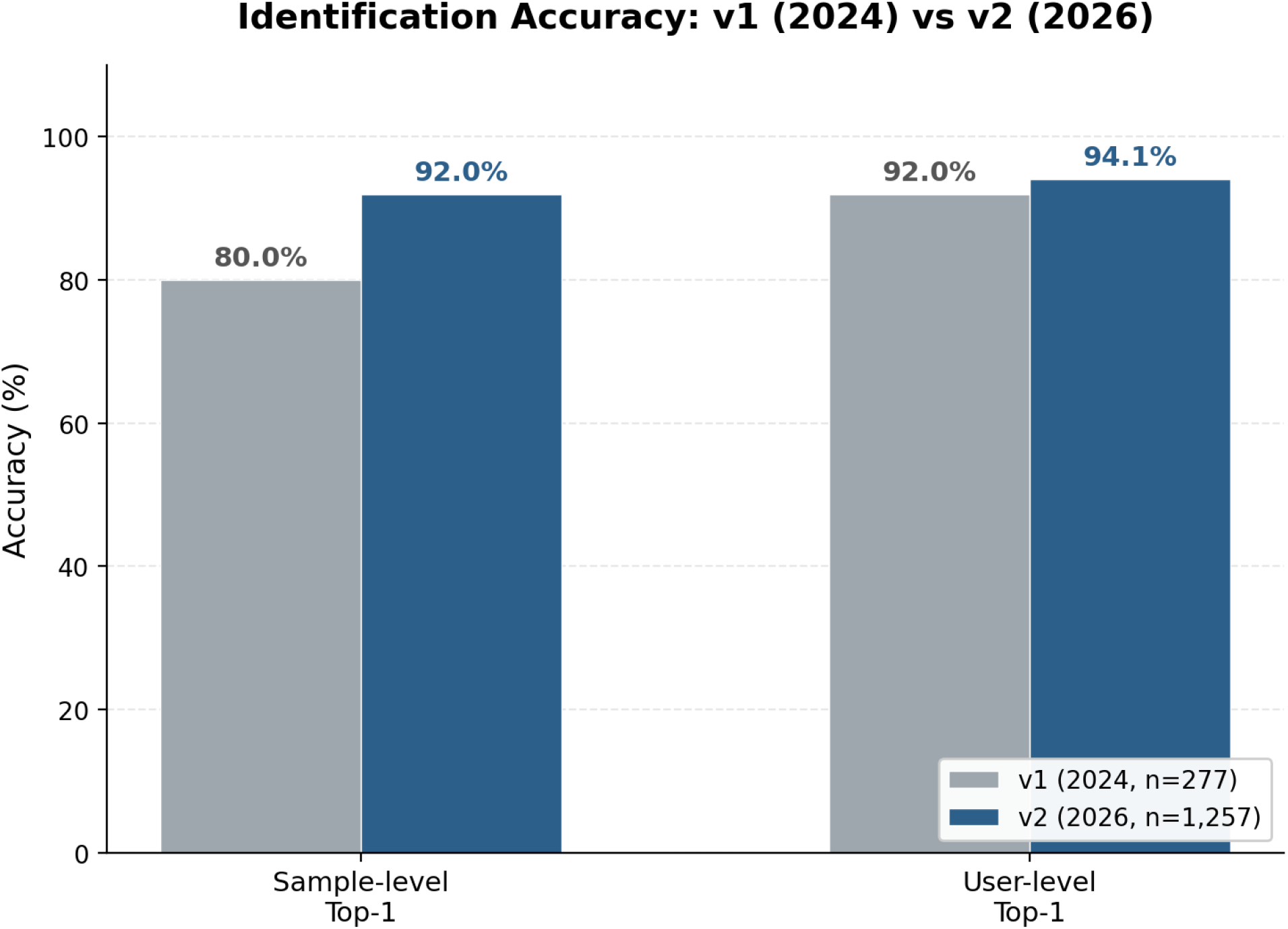
Accuracy comparison: v1 (2024) vs. v2 (this study) Left bars: sample-level Top-1 (v1: 80%, v2: 92.0% held-out). Right bars: user-level Top-1 (v1: 92% with 10 samples/user on 87 users, v2: 94.1% GroupKFold on 1,094 users).

**Figure 2.**
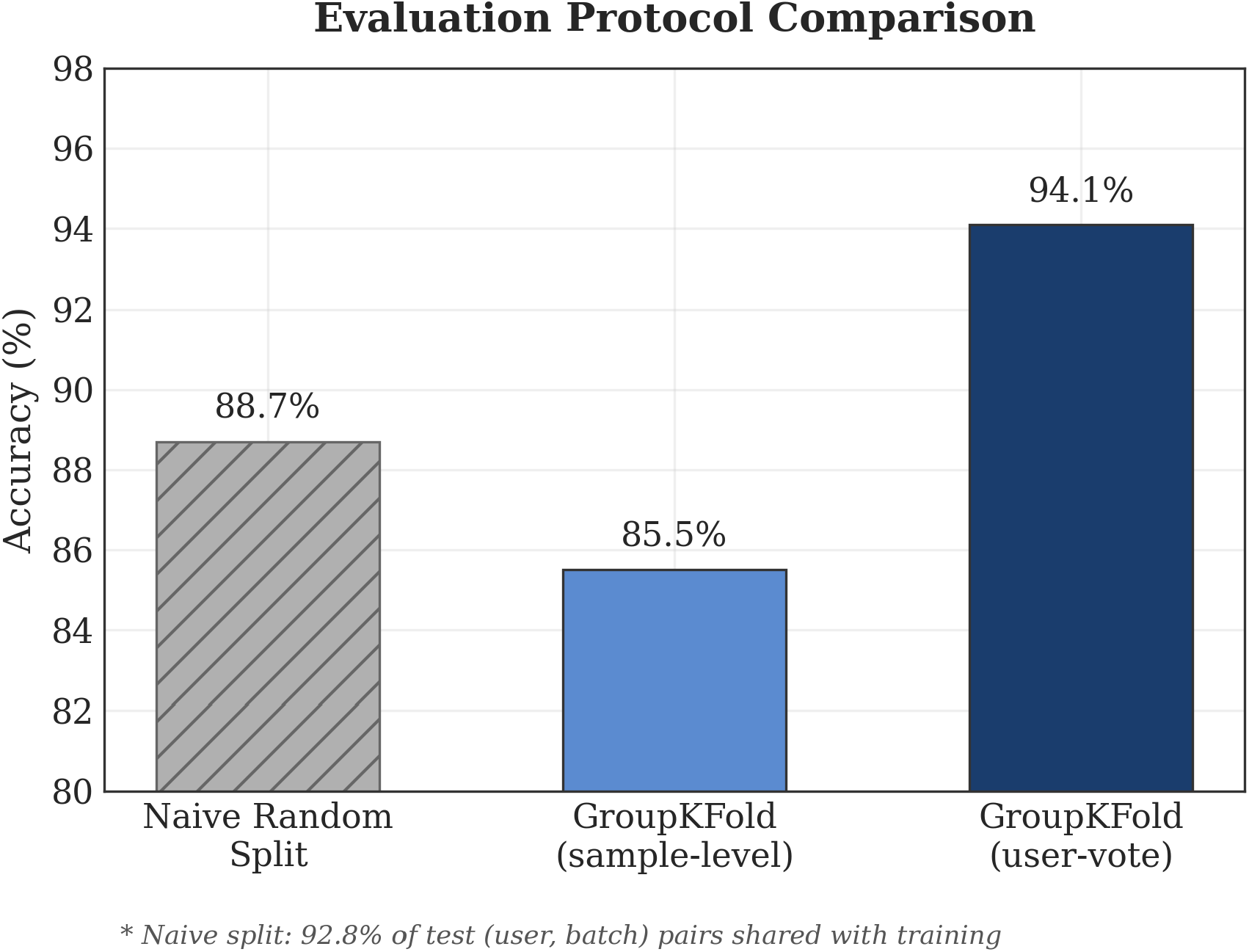
Identification accuracy under different evaluation protocols. Three bars shown: naive random split (88.7% sample-level, hatched, inflated by 92.8% batch leakage), GroupKFold sample-level (85.5%), and GroupKFold user-vote (94.1%). Both GroupKFold metrics ensure zero batch leakage.

**Figure 3.** Top discriminative metabolite classes by pathway. Color-coded by MSI annotation confidence: green (Level 1–2), orange (Level 3), red (contaminant). Removing all contaminants reduces held-out accuracy by only 0.5 percentage points.

**Figure 4.**
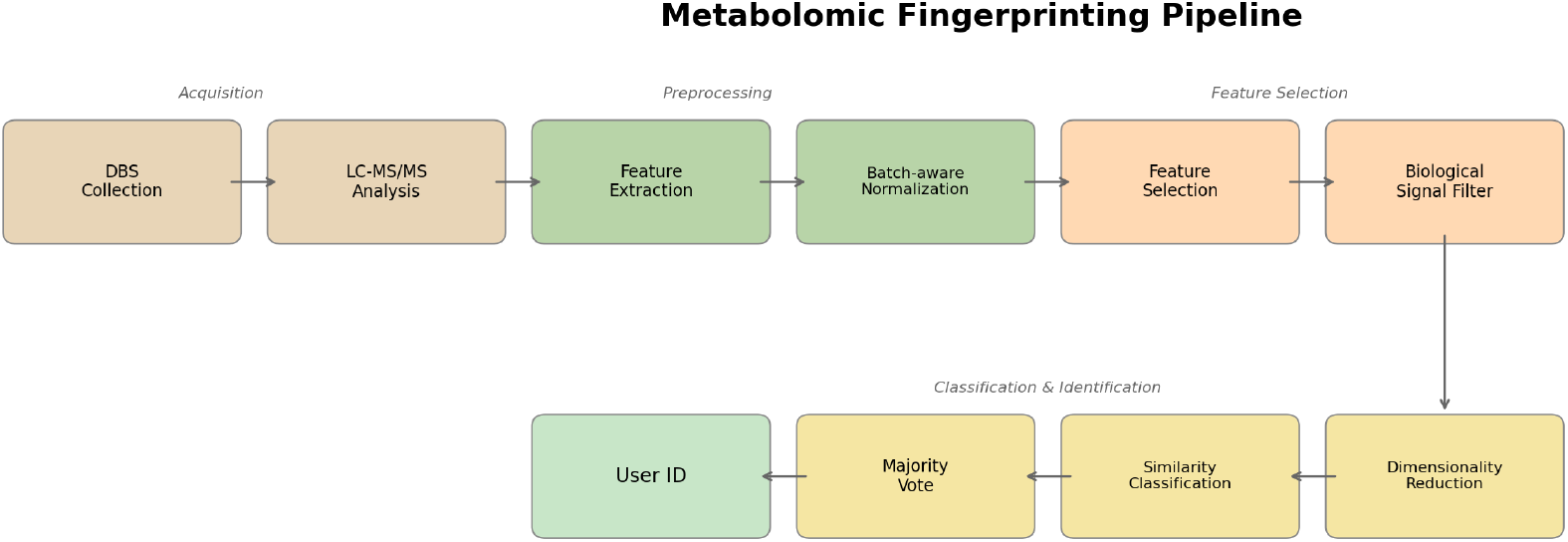
Pipeline overview: DBS collection to user identification. All pipeline steps are computed on training data only in the held-out validation.

